# Targeted sequence capture array for phylogenetics and population genomics in the Salicaceae

**DOI:** 10.1101/2020.05.08.084640

**Authors:** Brian J. Sanderson, Stephen P. DiFazio, Quentin C. Cronk, Tao Ma, Matthew S. Olson

**Affiliations:** Department of Biological Sciences, Texas Tech University, Lubbock, TX 79409-3131 USA; Department of Biology, West Virginia University, Morgantown, WV, 26506 USA; Department of Botany, University of British Columbia, Vancouver, BC, V6T 1Z4 Canada; Key Laboratory of Bio-Resource and Eco-Environment of Ministry of Education, College of Life Sciences, Sichuan University, Chengdu 610065, People’s Republic of China

**Keywords:** Phylogenetics, *Populus*, Salicaceae, *Salix*, targeted sequence capture

## Abstract

**Premise of the study:** The family Salicaceae has proved taxonomically challenging, especially in the genus *Salix*, which is speciose and features frequent hybridization and polyploidy. Past efforts to reconstruct the phylogeny with molecular barcodes have failed to resolve the species relationships of many sections of the genus.

**Methods:** We used the wealth of sequence data in the family to design sequence capture probes to target regions of 300-1200 base pairs of exonic regions of 972 genes.

**Results:** We recovered sequence data for nearly all of the targeted genes in three species of *Populus* and three species of *Salix*. We present a species tree, discuss concordance among gene trees, as well as some population genomic summary statistics for these loci.

**Conclusions:** Our sequence capture array has extremely high capture efficiency within the genera *Populus* and *Salix*, resulting in abundant phylogenetic information. Additionally, these loci show promise for population genomic studies.

## Introduction

Although the cost of whole-genome sequencing has continued to dramatically decrease over the past decade, the cost and complexity of whole-genome analyses still limit their utility and accessibility for answering evolutionary questions in novel taxa (Richards, 2018). However, a polished genome assembly is not necessary to address many questions. In this context, several methods have been developed to reduce the cost and effort required to obtain genomic information in novel species (McKain et al., 2018). The recent development of targeted sequence capture presents an affordable method for consistently isolating specific, long, phylogenetically informative regions in the taxa of interest (Gnirke et al., 2009; Mamanova et al., 2010; Hale et al., 2020). Targeted sequence capture uses biotinylated RNA baits to target prepared sequencing library fragments. The baited library fragments can then be pulled out of solution with streptavidin-coated magnetic beads to selectively enrich the fragments that contain loci of interest, while discarding the majority of library fragments that do not. This method offers many advantages over other methods of genome sequence partitioning, such as genome skimming and RAD-seq. It does not necessarily depend on a highly polished, annotated reference genome. Additionally, the same loci can be consistently sequenced at a high depth across individuals without requiring comprehensive, concurrent sequencing of all individuals (Mamanova et al., 2010; Grover et al., 2012; Jones and Good, 2016).

In this paper, we report on the design and implementation of a targeted sequence capture array to collect data for phylogenetic analysis within the Salicaceae, the plant family that includes poplars and willows. Understanding species relationships within this family, and in particular the genus *Salix*, has presented challenges to taxonomists as early as Linnaeus, who noted that “species of this genus are extremely difficult to clarify” (Linnaeus, 1753; Skvortsov, 1999). *Salix* species present challenges to classification due to their wide geographic ranges, hysteranthous phenology, extensive interspecific hybridization, polyploidy, and the lack of well-defined and variable flower characters for morphological circumscription of taxa (Raup, 1959; Skvortsov, 1999; Percy et al., 2014; Wang et al., 2020). Species of *Salix* exhibit holarctic distributions, and there are several classifications which differ among continents and are challenging to synthesize due to non-overlapping taxonomic treatment of species (Dickmann and Kuzovkina, 2014). Past efforts to reconstruct the phylogeny of *Salix* using nuclear AFLP and plastid barcode sequences have resulted in a lack of clearly resolved species relationships, especially in the subgenus *Vetrix* (Trybush et al., 2008; Percy et al., 2014). A more recent study using a supermatrix approach with RAD-seq data showed resolution within a subset of species of the subgenera *Vetrix* and *Chamaetia*, highlighting the potential of large-scale molecular data to resolve this phylogenetically challenging group (Wagner et al., 2018).

The utility of RAD-seq for collecting data for phylogeny, however, is limited by several issues. First, RAD-seq does not consistently screen homologous regions across species and across different experiments, which limits its utility for adding species to a phylogeny at a later time. Second, because RAD-seq assesses diversity in very short segments of the genome, the concatenation of this sort of data and supermatrix approaches are required for its use in phylogenetic analyses requires (de Queiroz and Gatesy, 2007), which does not allow separate exploration of gene and species phylogenies using super tree methods (Sanderson et al., 1998). Additionally, concatenation approaches are likely to exacerbate problems associated with maximum-likelihood methods for species with rapid diversification (Edwards et al., 2007; Edwards, 2009). Targeted sequence capture does not have these limitations, and thus may be a more appropriate genotyping platform for phylogenetics.

Species of *Populus* and *Salix* have been of great interest for the development of forestry and biofuel products, resulting in polished reference genomes for *P. trichocarpa, P. tremula, P. euphratica, S. purpurea* and *S. suchowensis*., as well as shallow resequencing data for many additional species (Tuskan et al., 2018). Our design strategy leveraged this abundance of existing genomic information to quantify polymorphism and the distribution of insertion-deletion events within and among species in order to maximize capture efficiency. Additionally, because we consistently target regions of exons we are able to characterize the nucleotide-site degeneracy with these data to quantify population genomic summary statistics. We demonstrate the utility of this resource for *Populus* and *Salix* species by presenting a fully resolved phylogenetic tree for six species and an outgroup, and by estimating the distribution of nucleotide diversity within species for our targeted genes.

## Methods

### Probe Design

Our goal was to identify regions that could be efficiently captured using RNA bait hybridization for diverse species across the family Salicaceae. The family Salicaceae is thought to have diverged from other clades approximately 92.5 Mya (Zhang et al., 2018b). Our primary focus was on the genera *Populus* and *Salix*, which diverged approximately 48 Mya, and the species *Idesia polycarpa* Maxim, which diverged from other clades approximately 56 Mya, which we use as an outgroup (Zhang et al., 2018b). Although we were interested in using these probes for phylogenetics with both *Populus* and *Salix* species, we focused on maximizing capture efficiency for the species in *Salix*, because the phylogeny for *Populus* is already much better resolved than that for *Salix* (Trybush et al., 2008; Wang et al., 2014, 2020; Percy et al., 2014; Liu et al., 2017). For this reason, the capture baits were designed to target regions in *Salix pupurea* that also would have high capture efficiency across the Salicaceae. The efficiency of RNA bait binding, and thus capture efficiency, is reduced as target regions diverge due to sequence polymorphism (Lemmon and Lemmon, 2013). To improve capture efficiency, we quantified sequence polymorphism among whole-genome resequencing data from a diverse array of *Populus* and *Salix* species (Table S1). The whole-genome short reads of the *Populus* and *Salix* species were aligned to the *Populus trichocarpa* genome assembly version 3 (Tuskan et al., 2006) using bwa mem v. 0.7.12 with default parameters (Li, 2013). We used the *P. trichocarpa* genome as our initial reference because it was the most polished and annotated genome in genus. Variable sites and insertion-deletion mutations (indels) were identified using samtools mpileup (Li, 2011), and read depth for the variant calls was quantified using vcftools (Danecek et al., 2011). Custom Python scripts were used to identify variant and indel frequencies for all exons in the *P. trichocarpa* genome annotation (scripts available at https://github.com/BrianSanderson/phylo-seq-cap; Sanderson, 2020).

Orthologs for our candidate loci in the in the *Salix purpurea* 94006 genome assembly version 1 (*Salix purpurea* v1.0, DOE-JGI; Carlson et al., 2017; Zhou et al., 2018) were identified using a list of orthologs between the *P. trichocarpa* and *S. purpurea* prepared using a tree-based approach by Phytozome v 12 (Goodstein et al., 2012). We further screened candidate regions to exclude high-similarity duplicated regions by accepting only loci with single BLAST (Camacho et al., 2009) hits against the highly contiguous assembly of *S. purpurea* 94006 version 5 (Zhou et al., 2020), which is less fragmented than the *S. purpurea* 94006 version 1. Genes from the Salicoid whole-genome duplication were identified using MCScanX (Wang et al., 2012), using default parameters and selected those segments for which the average *K*_*S*_ value for paralogous genes was between 0.2 and 0.8. Genes for which at least 600 base pairs (bp) of exon sequence contained 2-12% polymorphism and fewer than two indels were selected for probe design by Arbor Biosciences (Ann Arbor, MI, USA). Probes were designed with 50% overlap across the targeted regions, so that each nucleotide position would potentially be captured by two probes. Finally, to ensure that loci with high divergence across the family would be captured, we identified targets with less than 95% identity (based on BLAST results) between *S. purpurea* and *P. trichocarpa* and designed supplementary probes from orthologs of these genes in the *Idesia polycarpa* genome.

### Library Preparation and Sequence Capture

Libraries for two individuals from each *Populus balsamifera* L., *P. tremula* L., *P. mexicana* Wesmael., *Salix nigra* Marshall, *S. exigua* Nutt., and *S. phlebophylla* Andersson (Table S3) were prepared using the NEBNext Ultra II DNA Prep Kit following the published protocol for this kit (New England Biolabs, Ipswitch, MA, USA), and quantified using an Agilent Bioanalyzer 2100 DNA 1000 kit (Agilent Technologies, Santa Clara, CA, USA). Libraries were pooled at equimolar concentrations into two pools of six prior to probe hybridization following the Arbor Biosciences myBaits protocol v 3.0.1 and Hale et al. (2020). The hybridized samples were subsequently pooled at equimolar ratios and sequenced at the Texas Tech Center for Biotechnology and Genomics using a MiSeq with the Micro chemistry and 150 bp paired-end reads (Illumina, Inc., San Diego, CA, USA).

### Analysis of Sequence Capture Data

Read data was trimmed for primer sequences and low quality scores using Trimmomatic v. 0.36 (Bolger et al., 2014). The trimmed read data, as well as the whole-genome reads for *I. polycarpa*, were assembled into gene sequences using the HybPiper pipeline (Johnson et al., 2016). We estimated the depth of read coverage across all targeted genes as well as at off-target sites in R.

The assembled amino acid sequences were aligned with mafft v. 7.310 with the parameters – localpair and –maxiterate 1000 (Katoh and Standley, 2013), converted into codon-aligned nucleotide alignments with pal2nal v. 14 (Suyama et al., 2006), and trimmed for quality and large gaps with trimal v. 1.4.rev15 with the parameter -gt 0.5 (Capella-Gutiérrez et al., 2009).

HybPiper provides warnings for genes that have multiple competing assemblies that are within 80% of the length of the target region, because the alternate alignments may indicate that those genes have paralogous copies in the genome. We estimated phylogenetic relationships using the full set of gene sequences recovered from our sequence capture data, as well as a restricted set of putatively single copy genes, based on our *a priori* list of paralogs between *S. purpurea* and *P. tricocarpa*, and supplemented by the list of paralog warnings from HybPiper.

We estimated gene trees using RAxML v. 8.2.10, specifying a GTR*Γ* model of sequence evolution (Stamatakis, 2014). A set of 250 bootstrap replicates was generated for each gene tree. We used ASTRAL-III to infer the species tree from the RAxML gene trees (Zhang et al., 2018a; Rabiee et al., 2019). Because all nodes are weighted equally during quartet decomposition in ASTRAL-III, we used sumtrees in the Python package DendroPy v. 4.4.0 to collapse nodes with less than 33% bootstrap support values prior to species tree estimation (Sukumaran and Holder, 2010). A set of 100 multilocus bootstrap replicates was generated for the species tree. We used phyparts to determine the extent of congruence among gene trees for each node in the species tree (Smith et al., 2015). Cladograms representing the gene tree congruence and alternate topologies were plotted with the scripts phypartspiecharts.py and minority_report.py, written by Matt Johnson (scripts available at https://github.com/mossmatters/phyloscripts).

Finally, we used custom Python scripts to quantify nucleotide diversity at synonymous and non-synonymous sites between the individuals of the same species, as well as correlations in values of per-site nucleotide diversity between all species. The scripts described above as well as the full details of these analyses including are available in Jupyter notebooks at https://github.com/BrianSanderson/phylo-seq-cap (Sanderson, 2020).

## Results

### Sequence capture efficiency

The final capture kit targets 972 genes covered by 12,951 probes based on the *S. purpurea* reference, and an additional 7049 (redundant) probes based on the *I. polycarpa* genome that target genes with the highest divergence between *S. purpurea* and *P. trichocarpa* identified by Phytozome. This included an average of 680 ± 309 (mean ± sd) probes on each *S. purpurea* chromosome (Table S2), with an average of 1098 ± 489 (mean ± sd) bp of exon sequence per gene. Of the 972 target genes, 593 are putatively single copy based on our identification of paralogs in the *S. purpurea* genome assembly, 142 represent pairs of paralogs from the shared Salicoid whole-genome duplication (i.e. 71 pairs of genes), and 237 are genes that have known paralogs for which we were not able to design targets in this kit (i.e. each of these genes has one or more paralogs in the *S. purpurea* genome that is not targeted by probes). We included a total of 1219 genes in the target file used to assemble the capture data, which includes the 972 targeted genes as well as paralogous copies for which probes were not designed. Because the issues of paralogy become more complex when we add species other than *S. purpurea* and *P. trichocarpa*, we advise using the HybPiper warnings of multiple competing long assemblies to assess paralogy in novel species following guidance here https://github.com/mossmatters/HybPiper/wiki/Paralogs. The sequences of the capture probes as well as the target reference file are accessible at https://github.com/BrianSanderson/phylo-seq-cap (Sanderson, 2020). The sequence capture kit is available from Arbor Biosciences (Ref#170424-30 “Salicaceae”).

Sequence capture efficiency was high among the libraries. We recovered 805,820 ± 178,482 reads (mean ± sd) from our *Populus* and *Salix* target capture libraries, of which 86.7 ± 1.15% (mean ± sd) mapped to the target sequence reference (Table 1). An average of 94.48 ± 1.37% of targeted exon sequences were covered by ≥ 10 reads. The average read depth was 44.65 ± 1.61 for on-target sites, and 14.48 ± 2.10 for off-target sites (Table S4).

**Table 1.** Coverage summary statistics for sequence capture read data. For each library, values represent the number of reads in the sequenced library, the number of those reads that mapped to the reference file for the targeted genes, the proportion of mapped reads, the number of targeted genes (out of 972) that had read data mapped to them, and the number of genes that had 25%, 50%, 75%, and 100% of the targeted sequences covered with > 10X reads. Footnote: the I_polycarpa_WGS-2 data is from whole-genome sequencing data, rather than targeted sequence capture, and thus the low percent of read mapping reflects the lack of target enrichment (although the read coverage across targets was comparable to the sequence capture libraries, Table S3).

| **Name** | **Num.**<br>**Reads** | **Reads**<br>**Mapped** | **Prop.**<br>**Mapped** | **Genes**<br>**Mapped** | **Genes with**<br>**25% Seq** | **Genes with**<br>**50% Seq** | **Genes with**<br>**75% Seq** | **Genes with**<br>**100% Seq** |
| --- | --- | --- | --- | --- | --- | --- | --- | --- |
| I_polycarpa_WGS-2 | 223470714 | 1653494 | 0.007 | 971 | 970 | 966 | 944 | 123 |
| P_balsamifera_MGR-01 | 614093 | 523321 | 0.852 | 972 | 971 | 960 | 884 | 122 |
| P_balsamifera_MGR-04 | 769303 | 659712 | 0.858 | 972 | 972 | 965 | 915 | 145 |
| P_mexicana_PM3 | 843032 | 739728 | 0.878 | 972 | 972 | 964 | 917 | 140 |
| P_mexicana_PM5 | 880962 | 768927 | 0.873 | 972 | 972 | 967 | 913 | 142 |
| P_tremula_R01-01 | 749220 | 638002 | 0.852 | 971 | 970 | 960 | 907 | 134 |
| P_tremula_R04-01 | 634625 | 539805 | 0.851 | 971 | 969 | 956 | 876 | 122 |
| S_exigua_SE002 | 1139616 | 998698 | 0.876 | 969 | 969 | 966 | 937 | 229 |
| S_exigua_SE053 | 843120 | 741938 | 0.88 | 969 | 969 | 964 | 928 | 195 |
| S_nigra_SG037 | 1166615 | 1028635 | 0.882 | 971 | 971 | 967 | 932 | 205 |
| S_nigra_SG051 | 602649 | 524993 | 0.871 | 971 | 970 | 961 | 903 | 136 |
| S_phlebophylla_SP15M | 753628 | 651791 | 0.865 | 972 | 972 | 967 | 939 | 204 |
| S_phlebophylla_SP7F | 672975 | 581147 | 0.864 | 972 | 972 | 967 | 925 | 203 |

### Phylogenetics

The species tree estimated with putatively single copy genes correctly paired all individuals of the same species and revealed a fully resolved phylogeny for the *Populus* and *Salix* species with 100% multilocus bootstrap support for all nodes (Fig. 1A). At least 85% of gene trees support the topology of the species tree (Fig. 1B), with the exceptions of the bipartition that separates *P. balsamifera* and *P. tremula*, and the bipartition that separates *S. phlebophylla* from the other *Salix* species, which had dominant alternate topologies that were supported by a large number of gene trees (Figs. S1 and S2). The topology of the species tree estimated with the full set of genes and known paralogs was nearly identical to the tree estimated with only the putatively single copy genes. The major difference between these trees was evident in the bipartition separating *P. balsamifera* and *P. tremula*, where there were a large number of alternative topologies supported by small numbers of gene trees (the top 3 were supported by 13, 11, and 10 gene trees; Fig. S3).

**Figure 1.**
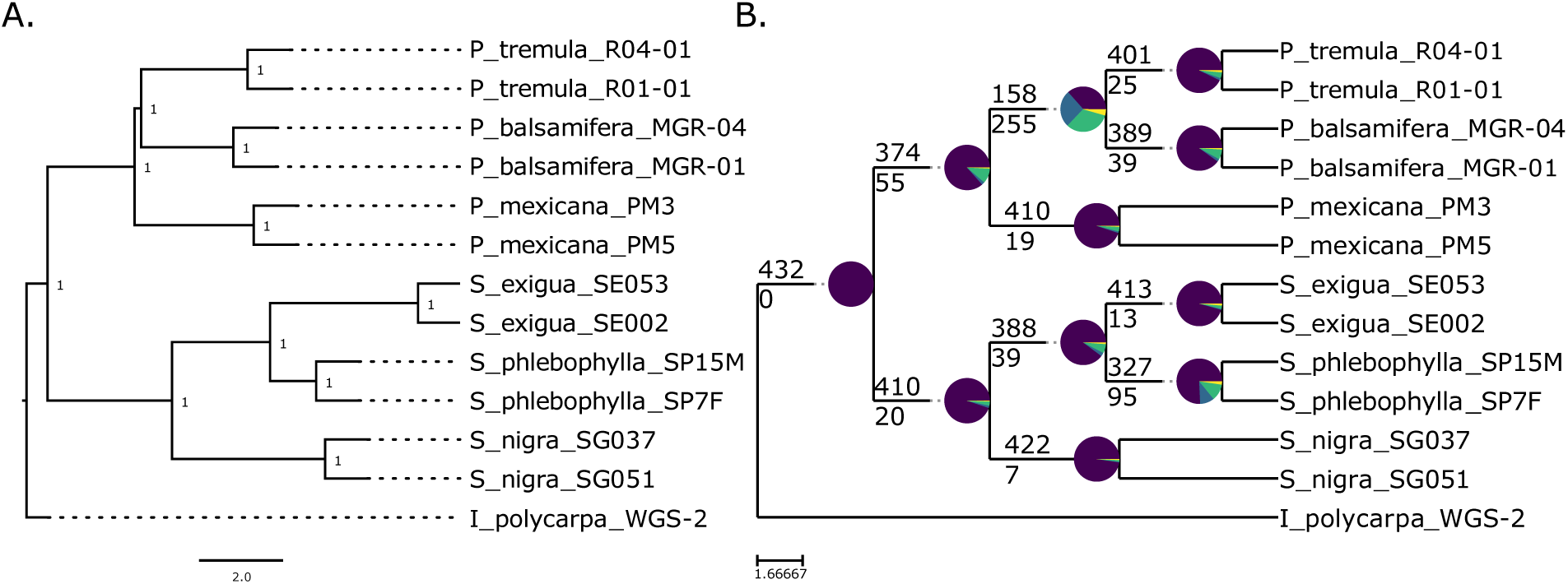
Species trees estimated for the 432 putatively single copy genes that did not have paralog warnings reported by HybPiper. **A)** Species tree generated by ASTRAL-III for the gene trees. Node values represent bootstrap support from 100 multilocus bootstrap replicates in ASTRAL-III. Branch lengths represent coalescent units. **B)** Cladogram showing the congruence of gene trees for all nodes in the ASTRAL-III species tree. The numbers above each node represent the number of gene trees that support the displayed bipartition, and numbers below the node represent the number of gene trees that support all alternate bipartitions. Purple wedges represent the proportion of gene trees that support the displayed bipartition. Blue wedges represent the proportion of gene trees that support a single alternative bipartition (see Figs S1 & S2). Green wedges represent the proportion of gene trees that have multiple conflicting bipartitions. Yellow wedges represent the proportion of gene trees that have no supported bipartition. Plotting code and its interpretation were provided by Matt Johnson (for more detail, see: https://github.com/mossmatters/MJPythonNotebooks/blob/master/PhyParts_PieCharts.ipynb)

### Population genomics

Patterns of nucleotide diversity, measured as Nei’s *π* (Nei and Li, 1979), varied among species, with the greatest variation at synonymous sites (Fig. S4; Table S5). *P. tremula* had the highest average values of *π* at both synonymous and non-synonymous sites (Fig. 2). The values of *π* among species were highly correlated for species within genus and exhibited lower correlations between genera (Fig. 3).

**Figure 2.**
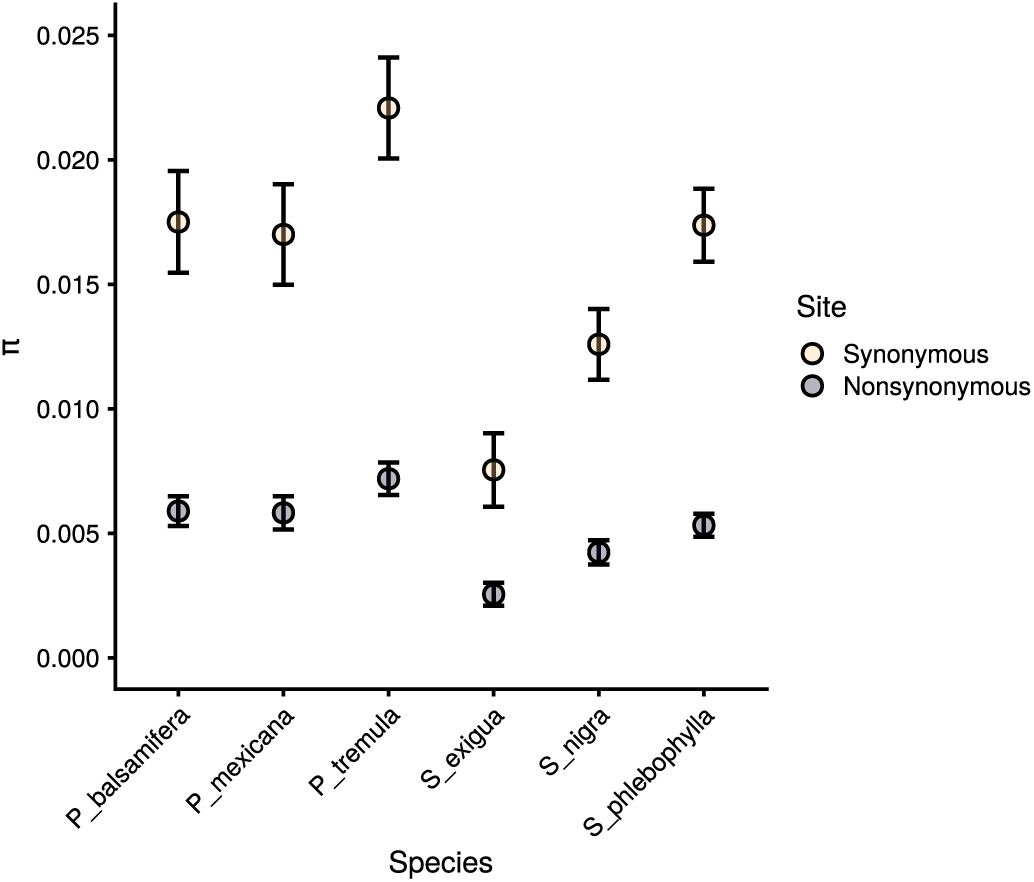
Means and 95% confidence intervals of values of nucleotide diversity (Nei’s π) within each species at synonymous (yellow) and nonsynonymous (purple) sites.

**Figure 3.**
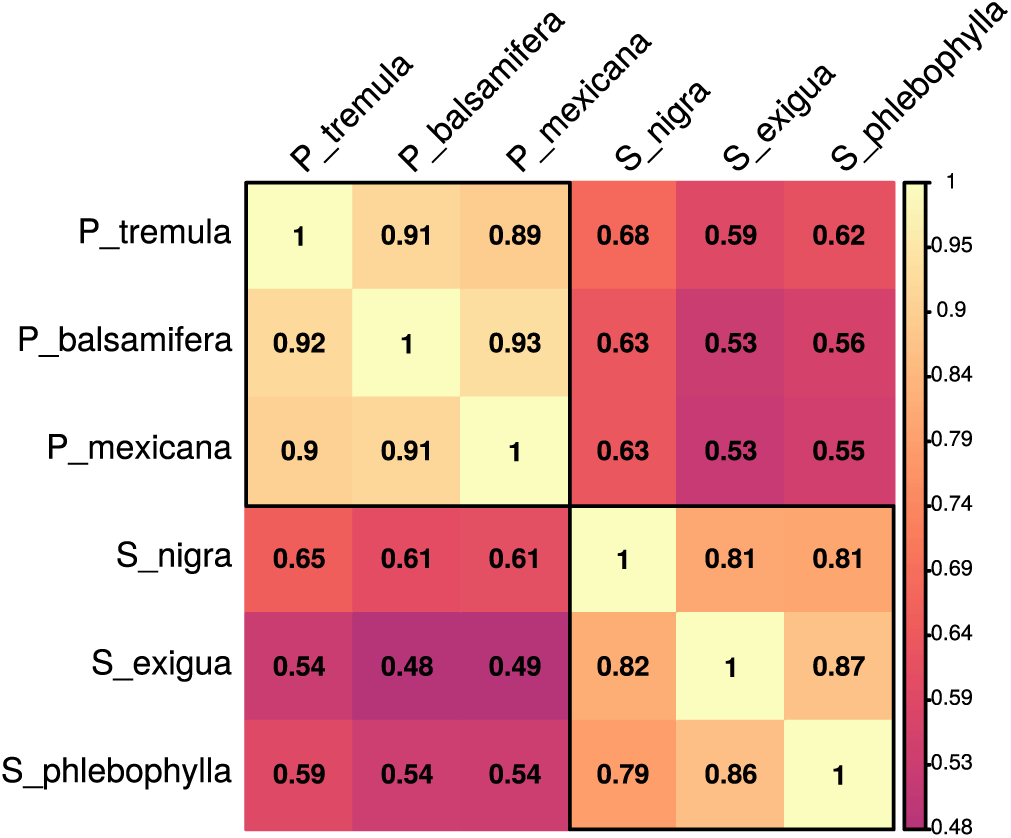
Pairwise correlation (Pearson’s r) of values of Nei’s π between all species. Values above the diagonal represent the correlation of π at synonymous sites, values below the diagonal represent non-synonymous sites. Black boxes represent within-genus comparisons.

## Discussion

The decreasing cost of obtaining genomic and transcriptomic sequence data holds great promise for unlocking our understanding of phylogenetic relationships and population genetic patterns within and among complex taxonomic groups. However, assembling complete genomes is still not a trivial task, and there exist relatively few polished plant reference genomes onto which genome skimming data can be mapped. Many methods have been developed to reduce the sequencing and analytical burdens associated with obtaining genome data. We believe that targeted sequence capture is one of the most promising contemporary methods of inexpensively generating genomic information.

The efficiency of our targeted sequence capture array was extremely high, which yielded abundant phylogenetic information for six species of *Populus* and *Salix*. Overall, the phylogeny was fully resolved and conformed to our general understanding the relationships among the taxa (Wu et al., 2015; Wang et al., 2020). One strength of the sequence capture approach is that it provides sufficiently long contiguous segments of gene sequences to assemble gene trees and it can overcome the problems introduced by concatenation of multiple gene regions with divergent histories (Edwards et al., 2007; Edwards, 2009). The super tree approach also allowed for the identification of alternative evolutionary histories that are supported by different regions of the genome, as often occurs during historical hybridization and introgression (Zhang et al., 2018a; Rabiee et al., 2019). Our species tree identified three alternative gene tree relationships among the three *Populus* species (Fig. S1). Previous studies have provided evidence of historical introgression among these species, including a history of chloroplast capture and hybridization between *P. mexicana* and species in the section *Tacamahaca* (including *P. balsamifera*; Wang et al., 2014, 2020; Liu et al., 2017). The second most supported alternative topology that we recovered placed *P. mexicana* and *P. tremula* as sister taxa, a pattern that does not support this hypothesis, likely due to incomplete lineage sorting (Wang et al., 2020). *P. tremula* likely has a greater long-term effective population size than *P. balsamifera* (Wang et al., 2016), and so coalescence times may be shorter on average in *P. balsamifera*. Among the *Salix* species, we identified three alternative gene tree relationships between the *S. phlebophylla* and *S. exigua* individuals, which may reflect the histories of rapid speciation and hybridization that have long vexed attempts at phylogenetic reconstruction in the genus *Salix* (Fig. S2; Trybush et al., 2008; Percy et al., 2014). Both of these patterns in *Populus* and *Salix* may be better understood once additional taxa are added to this phylogeny.

We have also shown that this sequence capture design can be applied to address questions related to population genomics in the Salicaceae. Many of the advantages of targeted sequence capture over competing methods are of particular relevance for population genomics studies, including specific knowledge of loci being sequenced, the ability to differentiate among synonymous, non-synonymous, intronic, and intergenic loci, and the ability to collect data on the same set of loci across different experiments, either within species or across species, for comparative studies. In particular, synonymous sites, especially four-fold synonymous sites, are among the fastest evolving regions of the genome and the sites within genic regions least influenced by selection (Wright and Andolfatto, 2008), and are thus among the best regions for estimating patterns of historical demography. Our estimates of nucleotide diversity are similar to those that have been previously reported for *P. balsamifera* and *P. tremula* using Sanger sequencing data (Ingvarsson, 2005; Olson et al., 2010) and whole-genome sequencing data (Wang et al., 2016). The high estimates of diversity in *S. phlebophylla* compared to the other two *Salix* species is curious and may result from a history with relatively little migration due to the absence of glaciation over a large portion of its Beringian distribution (Hultén, 1937).

The current study is based on a small sample size per species (n = 2), and so our ability to account for population structure or robustly perform population genomic inferences with these data is limited. Additionally, a potential limitation for using this sequence capture array for comparative population genomics is that we screened loci for a range of among-species variability between 2-12%, which excludes loci that exhibit extremely high or low values of nucleotide diversity. This may bias estimates of nucleotide diversity arising from these probes toward greater evenness. The ability to identify synonymous sites, which are the closest to neutral among all classes of sites (Wright and Andolfatto, 2008), should partially address this bias. Another feature of sequence capture data is the recovery of “off-target” sequences that result from the fact that the insert size of libraries is larger than the 120 bp bait length, and so regions upstream and downstream of the target will be sequenced as well. These regions may include intronic and intergenic regions, as well as exonic sequences that deviate from the constraints we used for our design. The results we report here only incorporate the “on-target” sites that we sequenced, but HybPiper implements methods to assemble intronic sequences as well. However, the potential effects of hitchhiking selection on synonymous site variation will likely remain apparent.

We also found that it was straightforward to integrate the targeted sequence capture data with whole-genome sequence data using the HybPiper pipeline by simply including the FASTQ files from whole-genome reads in the pipeline. This strategy was used to successfully incorporate whole-genome sequencing data from *Idesia polycarpa*, to act as our outgroup. The proportion of gene coverage as well as the read depth for the *I. polycarpa* data was similar to the sequence capture libraries (Table 1).

A whole genome duplication occurred prior to the divergence of *Salix* and *Populus*, and there are at least 8000 known paralog pairs in the *P. trichocarpa* reference genome (Tuskan et al., 2006). Genes with paralogous copies in the genome can complicate gene assemblies, because sequence data from both copies may alternately align to the same target sequence. We identified paralogous sequences in the *S. purpurea* genome assembly using MCScanX, and used that information to assist in the design the sequence capture array. The final array includes 593 putatively single copy genes, 142 pairs of paralogs, and 237 genes which have paralogs but for which we were not able to include both paralogs in the kit due our selection criteria. The target reference file we used to map the sequence capture data thus includes 1219 genes including the single copy and known paralogs from *S. purpurea*. In addition to this, HybPiper provides warnings for genes that have multiple competing alignments that cover the majority of the target sequence, which may indicate the presence of multiple paralogous copies in the genome (Johnson et al., 2016). This will be particularly useful because the genes that have maintained paralogous copies are likely to differ among species throughout the diversification of willows. We estimated evolutionary relationships using both the full set of 1219 single copy and known paralog genes, as well as a limited set of just single copy genes that did not report paralog warnings. The results from both analyses were nearly the same, but this will likely not be true for a more complex phylogenetic analysis that includes more than six species and an outgroup. For those more complex phylogenetic analyses, the ability to compare trees constructed with single copy genes with those using paralogous copies may provide crucial information for reconciling evolutionary relationships.

This sequence capture array will provide the community with an excellent resource to consistently sequence a set of variable regions of the genome for phylogenetic and population genomic investigations in the Salicaceae. The rate of read mapping and coverage of target genes was remarkably consistent across both genera, despite the fact that the taxa were selected to maximize sampling of phylogenetic diversity within each genus. The Salicaceae are important plants in the northern hemisphere both ecologically and economically and have been the subjects of numerous population genetics and genomics investigations of speciation, hybridization, introgression, selection, and historical population size and migration. This resource will allow phylogenetic and comparative population genomic studies to assess the same loci across different studies, which will allow us to build a worldwide diversity database and facilitate more precise comparative research questions. Our results demonstrate that the rate of gene capture is extremely high, such that it would be unnecessary to filter data and determine appropriate overlapping genotype thresholds, as is necessary with random genome partitioning methods such as RAD-seq.

## Acknowledgements

We thank M. Johnson for help with data analysis, P. Ingvarsson for help collecting *Populus tremula*, and J. Martinez, Instituto de Geología, UNAM, Hermosillo, Mexico for help collecting *Populus mexicana*. We thank J. Phillips for his work at Phytozome in identifying orthologs between *P. trichocarpa* and *S. purpurea*. Funding for this project was provided by the US National Science Foundation (1542509 and 1542599), Genome Canada (168BIO), and the National Natural Science Foundation of China (31561123001). The genetic material used to generate the genomic resources from AgCanBaP (Agriculture Canada Balsam Poplar) collection belong to Her Majesty the Queen in Right of Canada as represented by the Minister of Agriculture and Agri-Food. Agriculture and Agri-Food Canada retains complete ownership of the resources presented here.

## Author Contributions

S.P.D. and M.S.O. conceived the study. S.P.D., Q.C.C., T.M., and M.S.O. secured funding to support the project. B.J.S. and S.P.D. designed the sequence capture array. Q.C.C. and T.M. provided whole genome sequence data. B.J.S and M.S.O. prepared and sequenced the DNA samples, analyzed the data, interpreted the results, and wrote the manuscript. All authors edited drafts of the manuscript.

## Data Accessibility

Accession numbers for all sequence data used to design the sequence capture array are presented in Table S1. The raw reads of targeted sequence capture data from the six species of *Populus* and *Salix* are available on the NCBI sequence read archive under the BioProject accession number PRJNA627181. The raw reads of the *Idesia polycara* whole genome sequence data are available in the Genome Warehouse of the Beijing Institute of Genomics (BIG), under the accession number PRJCA002959. The sequences of the probes that were designed, all of the custom Python scripts that were used for this study, and the full details of analyses summarized in notebooks are available at https://github.com/BrianSanderson/phylo-seq-cap (Sanderson, 2020).

## Supporting Information

**Table S1**. Coverage summary statistics for whole-genome reads used to design sequence capture array. For each library, values represent the name of the sequenced individual, the number of reads in the sequenced library, the number of reads that mapped to the *Populus trichocarpa* v3 reference genome, the proportion of reads that mapped to the reference genome, and the mean and standard deviation of read depth.

**Table S2**. Distribution of probes across the *Salix purpurea* genome.

**Table S3**. Collection details for *Populus* and *Salix* species.

**Table S4**. Summary of read depth at on- and off-target sites. For each library values represent the 5%, 25%, 50%, 75%, and 99% quantiles of read depth, the maximum number of reads mapped to a site, and the mean and standard deviation of read depth.

**Table S5**. Nucleotide diversity expressed as Nei’s π for nonsynonymous and synonymous sites.

## Supplemental Figures

**Figure S1.**
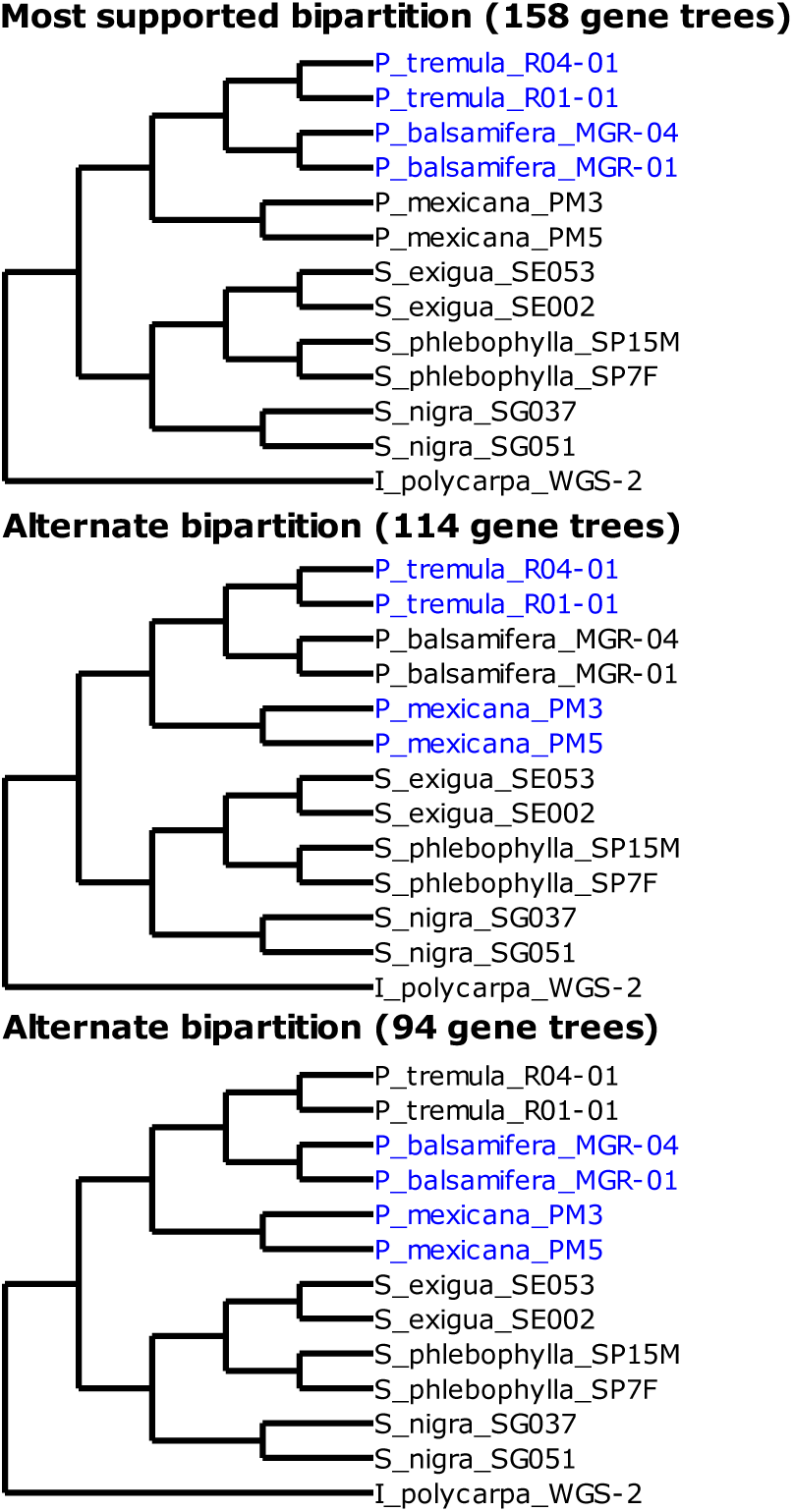
Alternate bipartitions for the three species of *Populus*, based on gene tree concordance. The cladogram in all three panels is that of the ASTRAL-III species tree (Figure 1), and the blue color represents the bipartition supported by the indicated number of gene trees in each panel. **A)** 158 gene trees support the displayed ASTRAL-III species tree topology. **B)** 114 gene trees support a bipartition that places *P. tremula* and *P. mexicana* together. **C)** 94 gene trees support a bipartition that places *P. balsamifera* and *P. mexicana* together.

**Figure S2.**
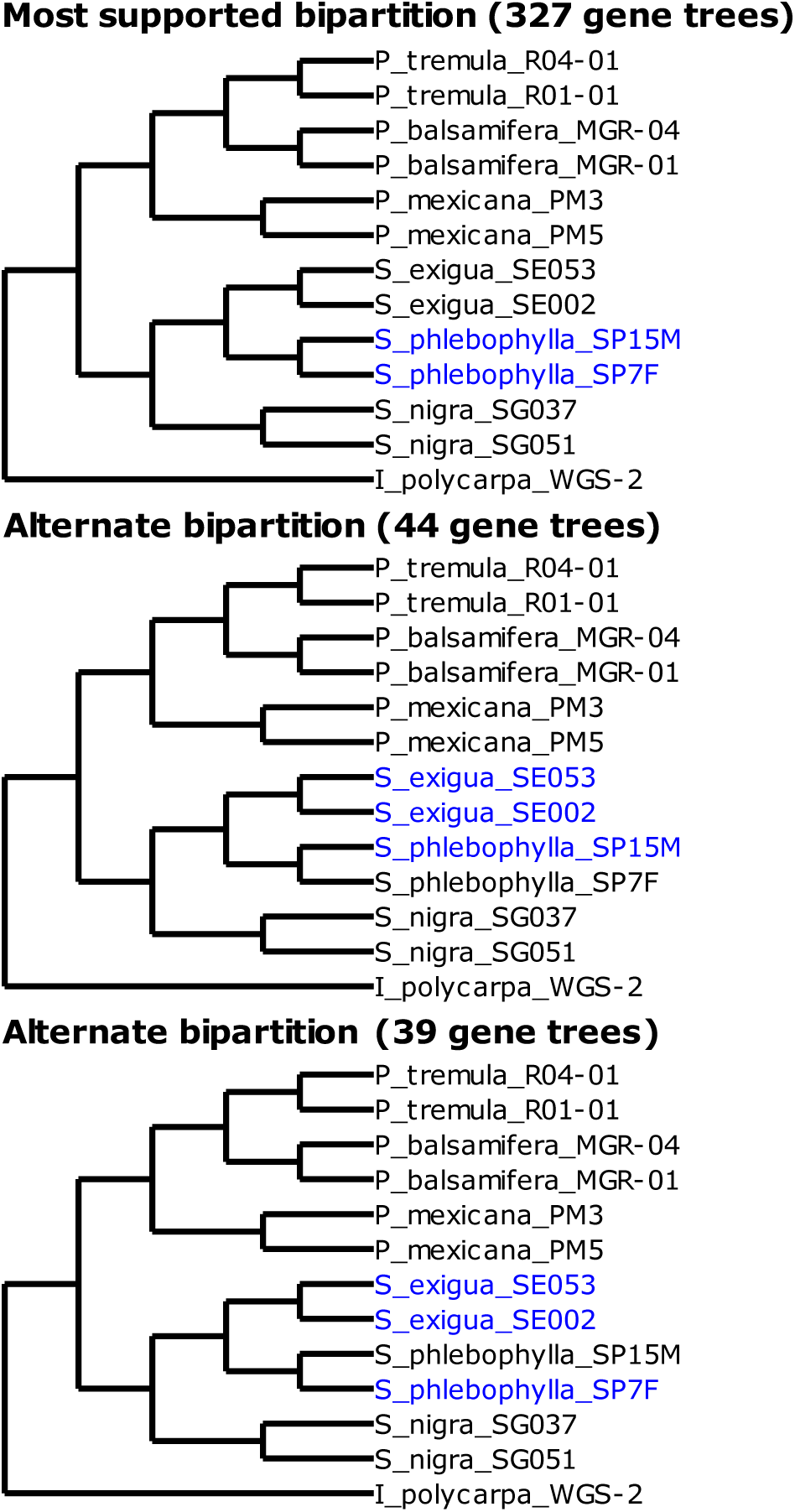
Alternate bipartitions for the three species of *Salix*, based on gene tree concordance. The cladogram in all three panels is that of the ASTRAL-III species tree (Figure 1), and the blue color represents the bipartition supported by the indicated number of gene trees in each panel. **A)** 327 gene trees support the displayed ASTRAL-III species tree topology. **B)** 44 gene trees and **C)** 39 gene trees place one of the *S. phlebophylla* individuals within *S. exigua*.

**Figure S3.**
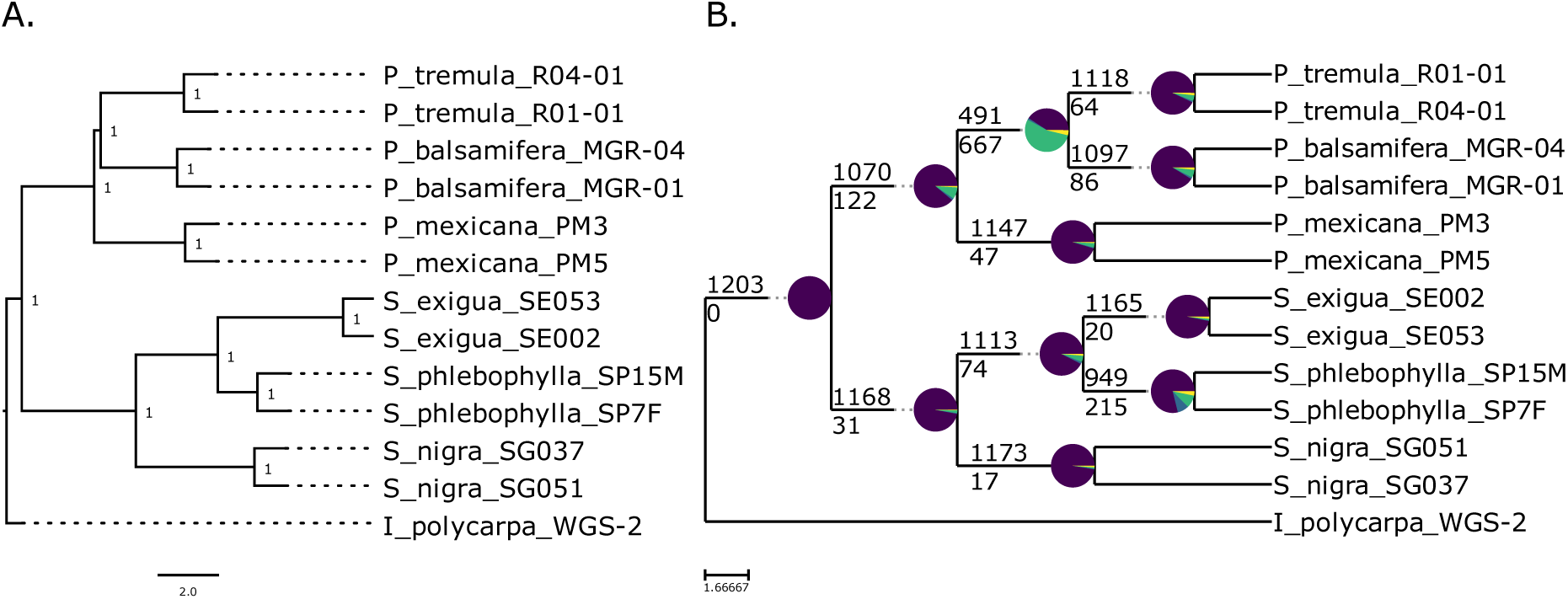
Species trees estimated for all genes and known paralogs. **A)** Species tree generated by ASTRAL-III for the gene trees. Node values represent bootstrap support from 100 multilocus bootstrap replicates in ASTRAL-III. Branch lengths represent coalescent units. **B)** Cladogram showing the congruence of gene trees for all nodes in the ASTRAL-III species tree. The numbers above each node represent the number of gene trees that support the displayed bipartition, and numbers below the node represent the number of gene trees that support all alternate bipartitions. Purple wedges represent the proportion of gene trees that support the displayed bipartition. Blue wedges represent the proportion of gene trees that support a single alternative bipartition. Green wedges represent the proportion of gene trees that have multiple conflicting bipartitions. Yellow wedges represent the proportion of gene trees that have no supported bipartition. Plotting code and its interpretation were provided by Matt Johnson (for more detail, see: https://github.com/mossmatters/MJPythonNotebooks/blob/master/PhyParts_PieCharts.ipynb)

**Figure S4.**
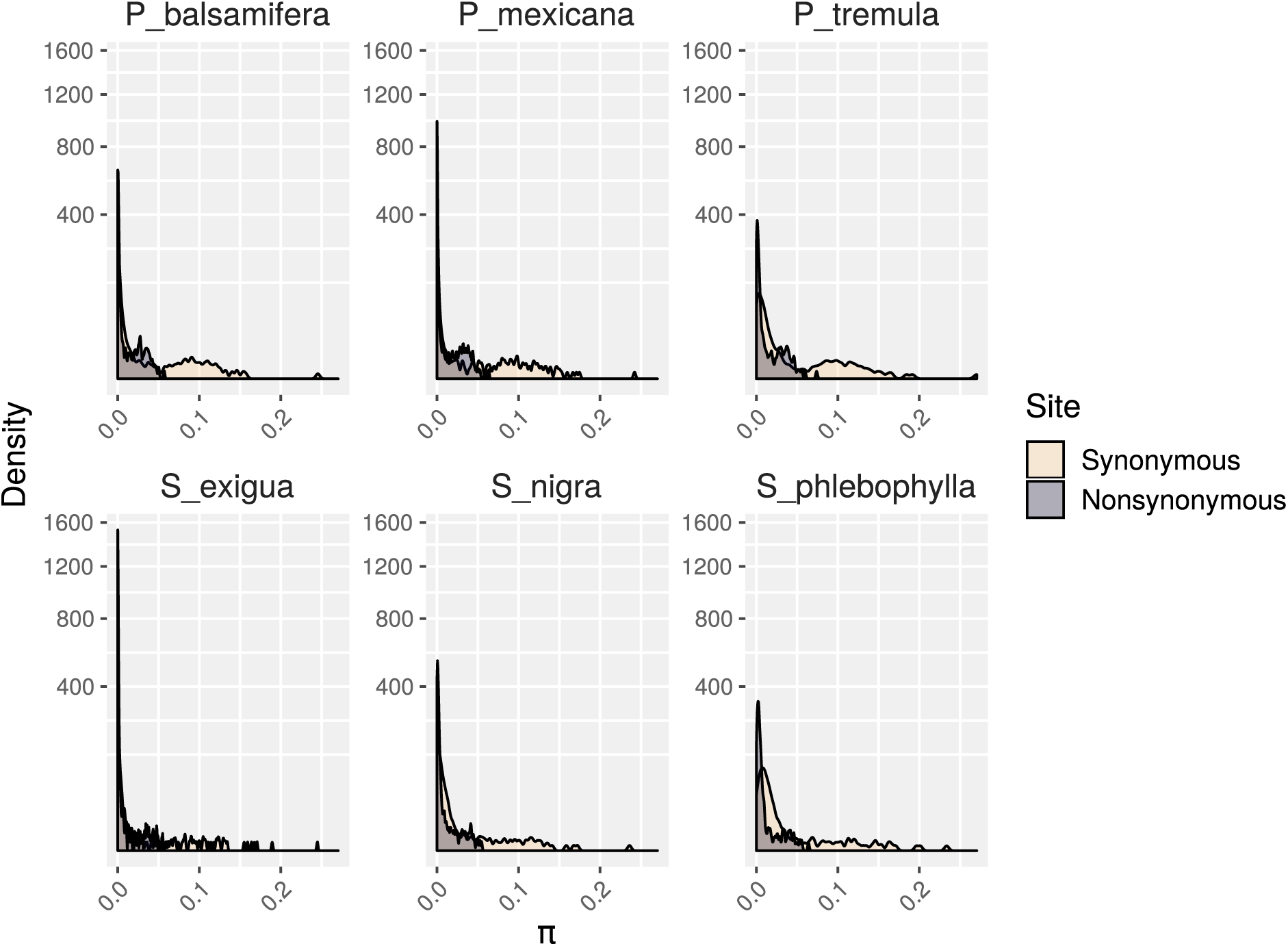
Distributions of values of nucleotide diversity (Nei’s π) within each species at synonymous (yellow) and nonsynonymous (purple) sites.

## Notes

### Competing Interest Statement

The authors have declared no competing interest.

### Summary of Updates

Light revisions mostly pertaining to a Zenodo release of data associated with this paper, as well as completion of the SRA accession numbers for raw read data in Table S1

## Literature Cited

Bolger, A.M., M. Lohse, and B. Usadel. 2014. Trimmomatic: A flexible trimmer for Illumina sequence data. Bioinformatics 30: 2114–2120.

Camacho, C., G. Coulouris, V. Avagyan, N. Ma, J. Papadopoulos, K. Bealer, and T.L. Madden. 2009. BLAST+: Architecture and applications. BMC Bioinformatics 10: 421–421.

Capella-Gutiérrez, S., J.M. Silla-Martínez, and T. Gabaldón. 2009. trimAl: A tool for automated alignment trimming in large-scale phylogenetic analyses. Bioinformatics 25: 1972–1973.

Carlson, C.H., Y. Choi, A.P. Chan, M.J. Serapiglia, C.D. Town, and L.B. Smart. 2017. Dominance and Sexual Dimorphism Pervade the Salix purpurea L. Transcriptome. Genome Biology and Evolution 9: 2377–2394.

Danecek, P., A. Auton, G. Abecasis, C.A. Albers, E. Banks, M.A. Depristo, R.E. Handsaker, et al. 2011. The variant call format and VCFtools. Bioinformatics 27: 2156–2158.

De Queiroz, A., and J. Gatesy. 2007. The supermatrix approach to systematics. Trends in Ecology and Evolution 22: 34–41.

Dickmann, D.I., and J. Kuzovkina. 2014. Poplars and Willows of the World, With Emphasis on Silviculturally Important Species. In J. Isebrands, and J. Richardson [eds.], Poplars and Willows Trees for Society and the Environment, 8–91. The Food and Agriculture Organization of the United Nations, Rome, Italy.

DOE-JGI. Salix purpurea version 1.

Edwards, S.V. 2009. Is a new and general theory of molecular systematics emerging? Evolution 63: 1–19.

Edwards, S.V., L. Liu, and D.K. Pearl. 2007. High-resolution species trees without concatenation. Proceedings of the National Academy of Sciences 104: 5936–5941.

Gnirke, A., A. Melnikov, J.R. Maguire, P. Rogov, E.M. Leproust, W. Brockman, T.J. Fennell, et al. 2009. Solution hybrid selection with ultra-long oligonucleotides for massively parallel targeted sequencing. Nature Biotechnology 27: 182–189.

Goodstein, D.M., S. Shu, R. Howson, R. Neupane, R.D. Hayes, J. Fazo, T. Mitros, et al. 2012. Phytozome: A comparative platform for green plant genomics. Nucleic Acids Research 40: D1178–D1186.

Grover, C.E., A. Salmon, and J.F. Wendel. 2012. Targeted sequence capture as a powerful tool for evolutionary analysis. American Journal of Botany 99: 312–319.

Hale, H., E.M. Gardner, J. Viruel, L. Pokorny, and M.G. Johnson. 2020. Strategies for reducing per-sample costs in target capture sequencing for phylogenomics and population genomics in plants: Low-cost Hyb-Seq. Applications in Plant Sciences e11337.

Hultén, E. 1937. Outline of the history of arctic and boreal biota during the Quarternary period : Their evolution during and after the glacial period as indicated by the equiformal progressive areas of present plant species. Bokförlags aktiebolaget Thule, Stockholm, Sweden.

Ingvarsson, P.K. 2005. Nucleotide polymorphism and linkage disequilibrium within and among natural populations of European aspen (Populus tremula L., Salicaceae). Genetics 169: 945–953.

Johnson, M.G., E.M. Gardner, Y. Liu, R. Medina, B. Goffinet, A.J. Shaw, N.J.C. Zerega, and N.J. Wickett. 2016. HybPiper: Extracting Coding Sequence and Introns for Phylogenetics from High-Throughput Sequencing Reads Using Target Enrichment. Applications in Plant Sciences 4: 1600016–1600016.

Jones, M.R., and J.M. Good. 2016. Targeted capture in evolutionary and ecological genomics. Molecular Ecology 25: 185–202.

Katoh, K., and D.M. Standley. 2013. MAFFT Multiple Sequence Alignment Software Version 7: Improvements in Performance and Usability. Molecular Biology and Evolution 30: 772–780.

Lemmon, E.M., and A.R. Lemmon. 2013. High-Throughput Genomic Data in Systematics and Phylogenetics. Annual Review of Ecology, Evolution, and Systematics 44: 99–121.

Li, H. 2013. Aligning sequence reads, clone sequences and assembly contigs with BWA-MEM. 00: 1–3.

Li, H. 2011. A statistical framework for SNP calling, mutation discovery, association mapping and population genetical parameter estimation from sequencing data. Bioinformatics 27: 2987–2993.

Linnaeus, C. 1753. Species plantarum. Impensis Laurentii Salvii, Stockholm.

Liu, X., Z. Wang, W. Shao, Z. Ye, and J. Zhang. 2017. Phylogenetic and Taxonomic Status Analyses of the Abaso Section from Multiple Nuclear Genes and Plastid Fragments Reveal New Insights into the North America Origin of Populus (Salicaceae). Frontiers in Plant Science 7: 1–9.

Mamanova, L., A.J. Coffey, C.E. Scott, I. Kozarewa, E.H. Turner, A. Kumar, E. Howard, et al. 2010. Target-enrichment strategies for next-generation sequencing. Nature Methods 7: 111–118.

Mckain, M.R., M.G. Johnson, S. Uribe-Convers, D. Eaton, and Y. Yang. 2018. Practical considerations for plant phylogenomics. Applications in Plant Sciences 6: e1038–e1038.

Nei, M., and W.H. Li. 1979. Mathematical model for studying genetic variation in terms of restriction endonucleases. Proceedings of the National Academy of Sciences 76: 5269–5273.

Olson, M.S., A.L. Robertson, N. Takebayashi, S. Silim, W.R. Schroeder, and P. Tiffin. 2010. Nucleotide diversity and linkage disequilibrium in balsam poplar (Populus balsamifera). New Phytologist 186: 526–536.

Percy, D.M., G.W. Argus, Q.C. Cronk, A.J. Fazekas, P.R. Kesanakurti, K.S. Burgess, B.C. Husband, et al. 2014. Understanding the spectacular failure of DNA barcoding in willows (Salix): Does this result from a trans-specific selective sweep? Molecular Ecology 23: 4737–4756.

Rabiee, M., E. Sayyari, and S. Mirarab. 2019. Multi-allele species reconstruction using ASTRAL. Molecular Phylogenetics and Evolution 130: 286–296.

Raup, H.M. 1959. The willows of boreal Western America. Contributions from the Gray Herbarium of Harvard University 185: 3–95.

Richards, S. 2018. Full disclosure: Genome assembly is still hard. PLOS Biology 16: e2005894–e2005894.

Sanderson, B.J. 2020. BrianSanderson/phylo-seq-cap: Publication (Version v1.0). Zenodo. http://doi.org/10.5281/zenodo.3979562.

Sanderson, M.J., A. Purvis, and C. Henze. 1998. Phylogenetic supertrees: Assembling the trees of life. Trends in Ecology and Evolution 13: 105–109.

Skvortsov, A.K. 1999. Willows of Russia and Adjacent Countries. I. N. (. Kadis, G. R. Argus, and A. G. Zinovjev [eds.], University of Joensuu, Joensuu.

Smith, S.A., M.J. Moore, J.W. Brown, and Y. Yang. 2015. Analysis of phylogenomic datasets reveals conflict, concordance, and gene duplications with examples from animals and plants. BMC Evolutionary Biology 15: 150–150.

Stamatakis, A. 2014. RAxML version 8: A tool for phylogenetic analysis and post-analysis of large phylogenies. Bioinformatics 30: 1312–1313.

Sukumaran, J., and M.T. Holder. 2010. DendroPy: A Python library for phylogenetic computing. Bioinformatics 26: 1569–1571.

Suyama, M., D. Torrents, and P. Bork. 2006. PAL2NAL: Robust conversion of protein sequence alignments into the corresponding codon alignments. Nucleic Acids Research 34: W609–W612.

Trybush, S., Š. Jahodová, W. Macalpine, and A. Karp. 2008. A genetic study of a Salix germplasm resource reveals new insights into relationships among subgenera, sections, and species. Bioenergy Research 1: 67–79.

Tuskan, G.A., S.P. Difazio, S. Jansson, J. Bohlmann, I. Grigoriev, U. Hellsten, N. Putnam, et al. 2006. The Genome of Black Cottonwood, Populus trichocarpa (Torr. & Gray). Science 313: 1596–1604.

Tuskan, G.A., A.T. Groover, J. Schmutz, S.P. Difazio, A. Myburg, D. Grattapaglia, L.B. Smart, et al. 2018. Hardwood Tree Genomics: Unlocking Woody Plant Biology. Frontiers in Plant Science 9: 1799–1799.

Wagner, N.D., S. Gramlich, and E. HÖrandl. 2018. RAD sequencing resolved phylogenetic relationships in European shrub willows (Salix L. Subg. Chamaetia and subg. Vetrix) and revealed multiple evolution of dwarf shrubs. Ecology and Evolution 8: 8243–8255.

Wang, J., N.R. Street, D.G. Scofield, and P.K. Ingvarsson. 2016. Natural Selection and Recombination Rate Variation Shape Nucleotide Polymorphism Across the Genomes of Three Related Populus Species. Genetics 202: 1185–1200.

Wang, M., L. Zhang, Z. Zhang, M. Li, D. Wang, X. Zhang, Z. Xi, et al. 2020. Phylogenomics of the genus *Populus* reveals extensive interspecific gene flow and balancing selection. New Phytologist 225: 1370–1382.

Wang, Y., H. Tang, J.D. Debarry, X. Tan, J. Li, X. Wang, T.-H. Lee, et al. 2012. MCScanX: A toolkit for detection and evolutionary analysis of gene synteny and collinearity. Nucleic Acids Research 40: e49–e49.

Wang, Z., S. Du, S. Dayanandan, D. Wang, Y. Zeng, and J. Zhang. 2014. Phylogeny reconstruction and hybrid analysis of populus (Salicaceae) based on nucleotide sequences of multiple single-copy nuclear genes and plastid fragments. PLoS ONE 9:.

Wright, S.I., and P. Andolfatto. 2008. The Impact of Natural Selection on the Genome: Emerging Patterns in Drosophila and Arabidopsis. Annual Review of Ecology, Evolution, and Systematics 39: 193–213.

Wu, J., T. Nyman, D.-C. Wang, G.W. Argus, Y.-P. Yang, and J.-H. Chen. 2015. Phylogeny of Salix subgenus Salix s.l. (Salicaceae): Delimitation, biogeography, and reticulate evolution. BMC Evolutionary Biology 15: 31.

Zhang, C., M. Rabiee, E. Sayyari, and S. Mirarab. 2018a. ASTRAL-III: Polynomial time species tree reconstruction from partially resolved gene trees. BMC Bioinformatics 19: 153–153.

Zhang, L., Z. Xi, M. Wang, X. Guo, and T. Ma. 2018b. Plastome phylogeny and lineage diversification of Salicaceae with focus on poplars and willows. Ecology and Evolution 8: 7817–7823.

Zhou, R., D. Macaya-Sanz, C.H. Carlson, J. Schmutz, J.W. Jenkins, D. Kudrna, A. Sharma, et al. 2020. A willow sex chromosome reveals convergent evolution of complex palindromic repeats. Genome Biology 21: 38–38.

Zhou, R., D. Macaya-Sanz, E. Rodgers-Melnick, C.H. Carlson, F.E. Gouker, L.M. Evans, J. Schmutz, et al. 2018. Characterization of a large sex determination region in Salix purpurea L. (Salicaceae). Molecular Genetics and Genomics 293: 1437–1452.

